# Spatial analysis of tumor infiltrating lymphocytes based on deep learning using histopathology image to predict progression-free survival in colorectal cancer

**DOI:** 10.1101/2021.04.24.441275

**Authors:** Hongming Xu, Yoon Jin Cha, Jean R. Clemenceau, Jinhwan Choi, Sung Hak Lee, Jeonghyun Kang, Tae Hyun Hwang

**Affiliations:** Department of Quantitative Health Sciences, Cleveland Clinic, Cleveland, OH 44195, USA; School of Biomedical Engineering, Faculty of Electronic Information and Electrical Engineering, Dalian University of Technology, Dalian, 116024, China; Department of Pathology, Gangnam Severance Hospital, Yonsei University College of Medicine, Seoul, 06273 Republic of Korea; Department of Hospital Pathology, Seoul St. Mary’s Hospital, College of Medicine, The Catholic University of Korea, Seoul, 06591 Republic of Korea; Department of Surgery, Gangnam Severance Hospital, Yonsei University College of Medicine, Seoul, 06273 Republic of Korea

**Keywords:** colorectal cancer, tumor infiltrating lymphocytes, deep learning, prognosis, whole slide image

## Abstract

**Purpose:** This study aimed to explore the prognostic impact of spatial distribution of tumor infiltrating lymphocytes (TILs) quantified by deep learning (DL) approaches based on digitalized whole slide images stained with hematoxylin and eosin in patients with colorectal cancer (CRC).

**Methods:** The prognostic impact of spatial distributions of TILs in patients with CRC was explored in the Yonsei cohort (n=180) and validated in the TCGA cohort (n=268). Concurrently, two experienced pathologists manually measured TILs at the most invasive margin as 0-3 by the Klintrup-Mäkinen (KM) grading method and compared to DL approaches. Interobserver agreement for TILs was measured using Cohen’s kappa coefficient.

**Results:** On multivariate analysis of spatial TILs features derived by DL approaches and clinicopathological variables including tumor stage, Microsatellite instability, and KRAS mutations, TILs densities within 200 μm of the invasive margin (f_im200) was remained as the most significant prognostic factor for progression-free survival (PFS) (HR 0.004 [95% CI, 0.0001-0.1502], *p*=.002) in the Yonsei cohort. On multivariate analysis using the TCGA dataset, f_im200 retained prognostic significance for PFS (HR 0.031, [95% CI 0.001-0.645], *p*=.024). Interobserver agreement of manual KM grading based on Cohen’s kappa coefficient was insignificant in the Yonsei (κ=.109) and the TCGA (κ=.121), respectively. The survival analysis based on KM grading showed statistically significant different PFS from the TCGA cohort, but not the Yonsei cohort.

**Conclusions:** Automatic quantification of TILs at the invasive margin based on DL approaches showed a prognostic utility to predict PFS, and could provide robust and reproducible TILs density measurement in patients with CRC.

**Data and Code Availability:** Source code and data used for this study is available at the following link: https://github.com/hwanglab/TILs_Analysis

## Introduction

Standard treatment of colorectal cancer (CRC) included curative intent surgical resection followed by postoperative selective chemotherapy with or without radiotherapy.(1) In patients with unresectable CRC, chemotherapy is the mainstream of treatment option to sustain the survival durations or to convert into resectable status. Postoperative chemotherapy has its own role to decrease recurrence, however, its adoption is solely dependent on postoperative staging under current guidelines.(1) Personalized treatment is demanding, because of different survival outcomes in patients with CRC belonged to the same stages.

The presence of tumor-infiltrating lymphocytes (TILs) is increasingly recognized as an important biomarker in multiple cancer types.(2–5) Previous studies using immunohistochemical (IHC) staining of various T-cell markers suggested that densities of TILs in the tumor microenvironment are associated with survival outcomes in patients with CRC.(2) Particularly, the Immunoscore that quantifies TILs densities and spatial distributions showed a high prognostic value in patients with CRC, which allows a more precise stratification of patient prognosis.(6) However, application of the Immunoscore requires IHC staining and expert interpretation, which could incur a substantial cost and specialized facilities.(2) Few studies measuring TILs using hematoxylin and eosin (H&E) stained slides also demonstrated its positive role as prognostic indicators in patients with CRC.(4, 7–11) Nevertheless, standardizing methods to quantify them are known to be labor-intensive and pathologist-dependent.(7) Although assessing TILs is considered to be clinically important, the TILs has been limitedly used as a prognostic biomarker due to additional requirements of IHC staining as well as substantial lack of standardization of measurement.(6, 12)

With recent advances in digital pathology and artificial intelligence (AI) (i.e., deep learning algorithm), there have been increasing interests in developing automated methods for TILs quantification and analysis from pathology slides. Saltz and colleagues used deep learning approaches to detect TILs in H&E stained whole slide images (WSIs), and showed that spatial TILs distribution within tumors was linked to patient survival across different tumor types.(13) Corredor and colleagues performed H&E stained tissue microarray slides based analysis to identify TIL spatial distribution and its co-localization with cancer cell nuclei to predict likelihood of recurrence in early-stage non-small cell lung cancer.(12) Recent study showed that the TILs densities present in tumor regions detected by the QuPath® software (14) in H&E stained pathology images could be used to predict overall survival in patients with melanoma.(15) Although recent studies suggest that TILs-related variables extracted by computational pathology could potentially predict patient clinical outcomes across different cancer types, the quantitative analysis of TILs according to spatial distribution using H&E stained WSIs has been limitedly investigated in patients with CRC.

In this study, we aimed to investigate whether automated quantification of spatial distribution of TILs present in tumor invasive margins based on deep learning approaches utilizing H&E stained WSIs could predict progression-free survival (PFS) outcomes in patients with CRC. In addition, we also sought to compare the Deep Learning based TILs density measurement (DeepTILs) in terms of their ability to predict prognosis in these patients with CRC against the manual scoring of TILs at the deepest invasive area by pathologists.

## Materials and Methods

### Datasets

Two independent cohorts of H&E stained WSIs from 448 colorectal cancer patients were included in this study. The included cohorts are represented by the Yonsei (n=180) and TCGA (n=268) respectively. The Yonsei cohort consists of 180 diagnostic WSIs and corresponding clinical information from stage II or III colon cancer patients who underwent curative resection followed by FOLFOX (5-Fluorouracil, leucovorin, and oxaliplatin) chemotherapy between September 2005 and January 2014, which data were collected in Gangnam Severance Hospital, Yonsei University College of Medicine, Republic of Korea. The TCGA cohort were composed of 268 CRC patients whose clinical information and WSIs were downloaded from The Cancer Genome Atlas (TCGA) data portal (https://portal.gdc.cancer.gov/).

The WSIs from Yonsei dataset (.mrxs) were generated by using Pannoramic® 250 Flash III scanner (3DHISTECH, Hungary) with pixel resolution of 0.2428um/pixel. The TCGA pathology slides were generated and uploaded by many different institutions, where images (.svs) were scanned by using the Aperio Scanscope CS scanner with pixel resolution of 0.2527um/pixel.

Patients who had post-operative WSIs and available clinicopathologic and follow up data were included. In case of TCGA dataset, patients with very poor quality of slides, absence of survival status or duration, and patients whose survival time was denotes as 0 month were excluded. When the pathologists evaluated the manual scoring of TILs, they found that some slides did not include the invasive margins and these were excluded at the stage in measuring the agreement rating and further comparison processing between human scoring versus AI based scoring (Supplementary Figure 1).

For each patient, the following outcomes were collected if available: sex, age (years), American Society of Anesthesiologists (ASA) classification, body mass index (BMI) (kg/m^2^), carcinoembryonic antigen (CEA) (ng/mL), tumor location, complications, histologic grade, lymphovascular invasion (LVI), total retrieved lymph node numbers, stage, microsatellite instability (MSI) and KRAS mutation status.

This study was conducted after approval of the Institutional review board of the Gangnam Severance Hospital, Yonsei University College of Medicine (Seoul, Republic of Korea) (approval no. 3-2020-0076). The need for informed consent was waived for this retrospective study.

### Development of Deep Learning based TILs density measurement (DeepTILs) using H&E WSIs

The developmental process of DeepTILs was consisted of four main modules: tumor detector, TILs detector, automatic quantification of TILs and statistical and survival analysis (Figure 1). The details are described in the following.

**Figure 1.**
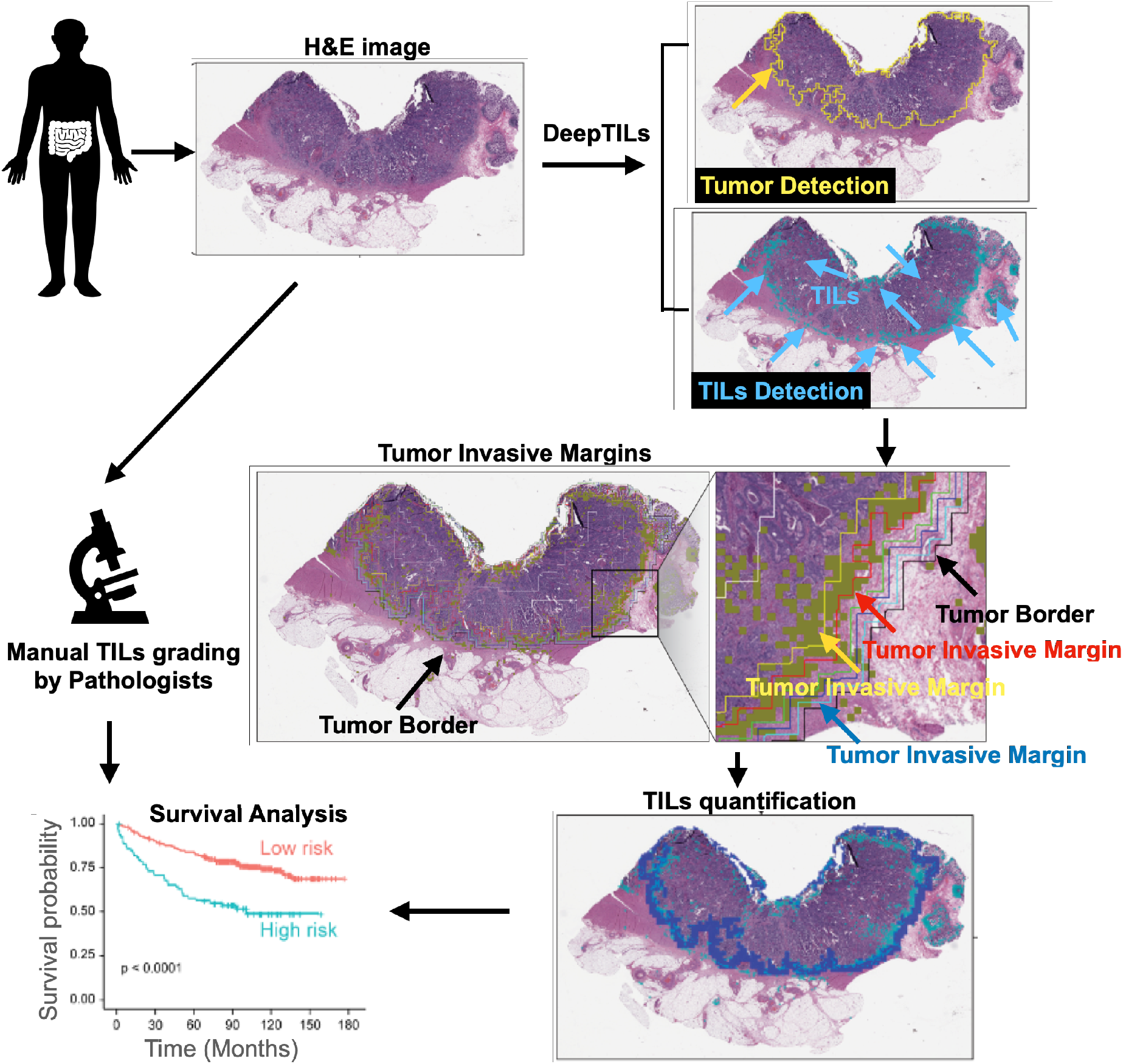
Overview of our approach. We first detect tumor region and TILs based on H&E whole slide image by Deep Learning approaches. Then quantify TILs across and within tumors. We used spatial distributions of TILs densities at tumor invasive margins and tumor core to identify patient subgroups with distinct progression free survival (PFS) outcome.

#### Tumor detector

To develop an automatic tumor detector module, we used the Resnet 18 deep learning model with a public dataset containing 11,977 image patches (256μm edge length per image) of H&E stained histological samples of human CRC.(16, 17) Regions in these images were manually annotated into three classes: tumor tissue, loose non-tumor tissue (i.e., adipose tissue and mucus), and dense non-tumor tissue (i.e., stroma and muscle). To train and test tumor detector, we randomly divided the whole dataset into training (80%), validation (10%) and testing (10%) sets. We then trained the Resnet18 deep learning model to distinguish tumor and non-tumor image patches. Image augmentations including random flipping and color jittering were applied along with training. We trained the model by freezing different percentile of trainable layers, and using different parameter configurations in terms of optimizer, batch size and learning rate (Supplementary Table 1). We trained and tested 24 different tumor detector models. Testing accuracies of the 24 models with different configurations were described in supplementary Figure 2. The tumor detector model which was trained by fine-tuning all trainable layers of Resnet18 and using Adam optimizer with learning rate of 0.0001 and batch size of 64 provided the best test accuracy (100%). This best performing tumor detector model was applied to predict tumor regions for all WSIs from the Yonsei and TCGA cohorts. Specially, the WSI was divided into a set of non-overlapping image tiles (256 μm x 256 μm per tile) which were predicted as tumor or non-tumor tiles by using our best tumor detector. The predicted image tiles were then stitched together to form WSI-level tumor detection results which are indicated by red pixels in Supplementary Figure 3-A (tumor detection).

#### TILs detector

In parallel with tumor detection, we trained and applied deep learning model to detect TILs regions in WSI. To develop the TILs detector models, we trained three different deep learning models (e.g., resent18, resent34, and shufflenet) with a public dataset, where 43,440 image tiles were adopted.(13) The whole data set was randomly divided into three parts: training (80%), validation (10%) and testing (10%) sets. Image augmentations including random flipping and color jittering were applied along with training. We trained the models by freezing different percentile of trainable layers, and using different parameter configurations in terms of optimizer, batch size and learning rate (Supplementary Table 2). We trained and tested 144 different TILs detectors, and testing accuracies of 144 models with different configurations were described in Supplementary Figure 4. The TILs detector model which was trained by fine-tuning all trainable layers of Resnet18 and using Adam optimizer with learning rate of 0.0001 and batch size of 4 provided the best test accuracy (80.06%). This best TILs detector model was used to make TILs detections for all WSIs from the Yonsei and TCGA cohorts. Specially, the WSI was divided into a set of 112 μm x 112 μm image patches which were predicted as the probabilities belonging to TILs. The WSI-level TILs prediction was finally obtained by stitching tile-level predictions. Supplementary Figure 3-B (TILs detection) illustrates TILs detection example (e.g., yellow pixels).

#### Automatic quantification of TILs from H&E stained WSIs

Based on predicted tumor and TILs regions by the deep learning models, we quantified the density of TILs insides invasive margin (IM) and tumor regions, respectively. If there exist more than one detected tumor regions, the largest tumor region is selected. We used image morphological dilation operation to detect the tumor IM. Note that by adjusting the physical size of structuring elements, we can obtain different width of IM layers. The schematic spatial distribution of TILs was illustrated in Supplementary Figure 5-A. The red contour indicates the tumor border, while different width of IM layers are indicated by different color lines to the distance of 200, 300, 400 and 500 μm from the tumor border. The TILs densities are computed and denoted as “f_im200”, “f_im300”, “f_im400” and “f_im500” at each IM layer, respectively. For example, “f_im200” represents the TILs densities of 200 μm IM layer. In addition to TILs densities at IM layers, we further quantified TILs densities at tumor regions. The TILs density at the whole tumor region is computed as “f_wt”. The area that occupies 25% of the central part of the whole tumor region is defined as the tumor core and the TILs density in the tumor core is denoted as “f_tc”. We applied morphological erosion operation to obtain 200μm inverse IM layer and computed the corresponding TILs density as “f_inv200”. Finally, in the whole tumor region, area of inverse 200 IM layer was subtracted and the remaining area was defined as TC2, and the TILs density of this area was defined as “f_tc2” (Supplementary Table 3, Supplementary Figure 5).

### Manual scoring of TILs by pathologists using Klintrup-Mäkinen (KM) recommendation

Two board-certified pathologists, who had 9 and 10 years of experiences respectively, graded each patient’s TILs by the Klintrup-Mäkinen (KM) recommendation (KM grading),(10) of which the deepest area of the invasive margin of the tumor was assessed by using a 4-degree scale. A score 0 denoted no increase in inflammatory cells, 1 denoted a mild and patchy increase in inflammatory cells, 2 denoted a moderate and band like inflammatory infiltrate with some destruction of cancer cell islands, and 3 denoted a marked and florid cuplike inflammatory infiltrate with frequent destruction of cancer cell islands.(8, 9) Patients were dichotomized as KM-low (KM grading 0 and 1) and KM-high (KM grading 2 and 3), and survival outcome were compared between these two groups.

### Statistical analysis

All statistical analyses were performed using R version 3.6.3 (R-project, Institute for Statistics and Mathematics, Vienna, Austria). Patients’ clinicopathological characteristics were compared using the chisquare and t-tests for categorical and continuous variables, respectively. Correlation between TIL counts and clinical variables were assessed by Mann-Whitney U test for dichotomous values, Kruskal-Wallis tests for more than three group comparisons, and Spearman correlations for continuous parameters.

The kappa statistics was used to assess inter-observer agreement with respect to scoring and was interpreted according to the guidelines of Landis and Koch.(18) We used the following definition to interpret the kappa coefficients: a kappa (κ) value of equal to or less than 0.20 indicated insignificant agreement; values from 0.21-0.40, median agreement; 0.41-0.60, moderate agreement; 0.61-0.80, substantial agreement; and 0.81-1.00, almost perfect agreement.

PFS was calculated from the date of surgery until the date of recurrence detection, last follow-up, or death. The patients alive at the last follow-up were censored. The Kaplan–Meier method was used to construct survival curves and the log-rank test was used to compare survival rates between groups. Cox proportional hazards models were used to estimate the hazard ratios (HRs) and 95% confidence interval (CI). All variables with a *p*<0.1 on univariate analysis were entered for multivariate analysis with backward stepwise selection of variables. A two-sided *p*<.05 was considered statistically significant.

## Results

### Clinicopathologic characteristics of the patient

Among the included patients, 29 of 180 patients (16.1%) in Yonsei cohort, and 74 of 268 patients (27.6%) in TCGA cohort had recurrences. The median follow-up periods for patients was 89 months (interquartile range [IQR], 71–122 months) for the Yonsei cohort and 19.6 months (IQR, 12.4–32.9 months) for the TCGA cohort. Clinicopathological features of the included patients were presented in Table 1.

**Table 1.**
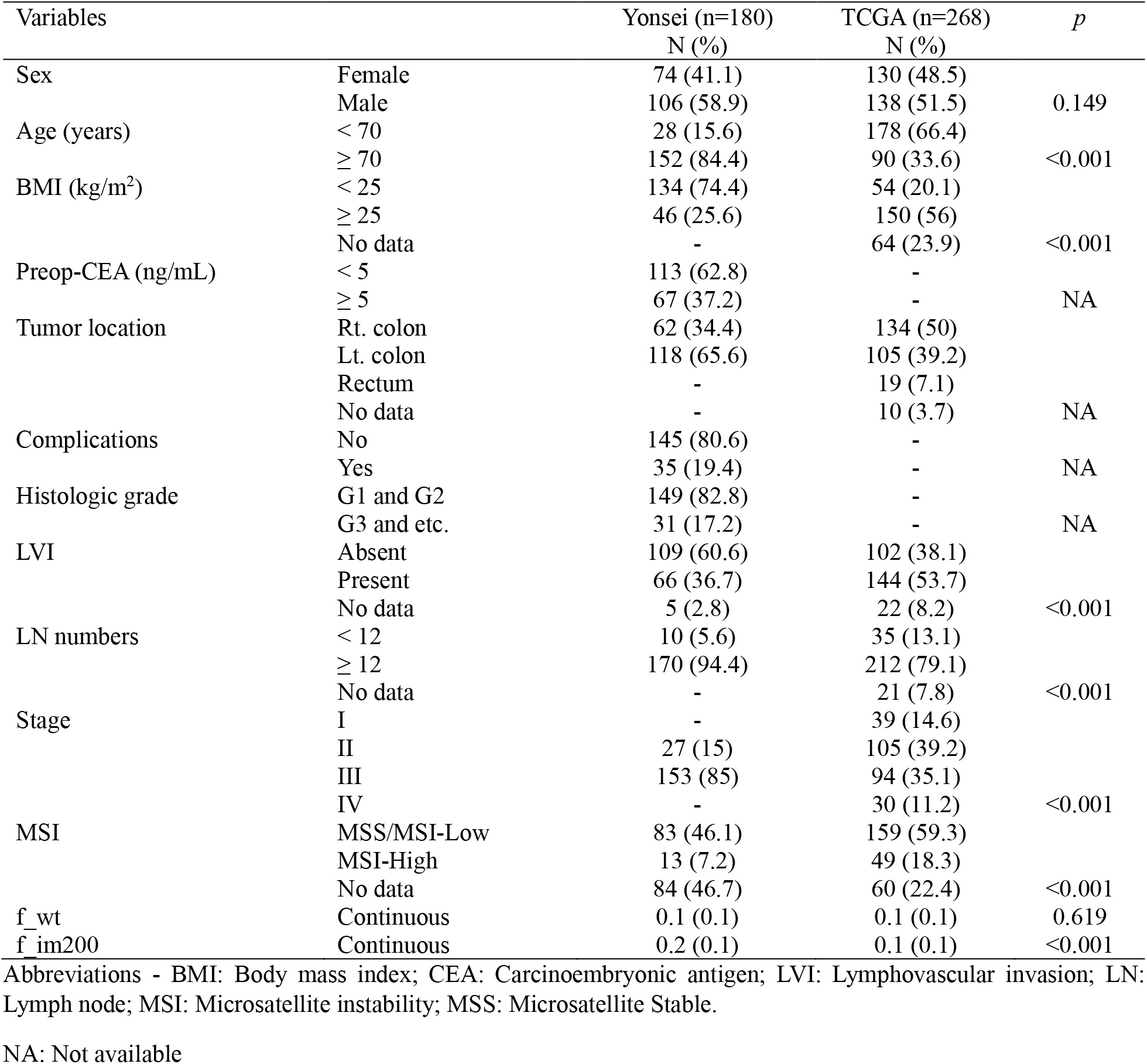
Patient characteristics of the Yonsei and TCGA dataset

### Prognostic evaluation of automated TILs features by DeepTILs from the Yonsei dataset

Composition of TILs according to spatial distribution was illustrated in the Supplementary Table 4. The median TILs density of each spatial distribution ranged from 0.0447 to 0.2002. Univariate Cox proportional-hazards model of PFS in the Yonsei cohort revealed that “f_im200”, “f_im300”, “f_im400”, “f_im500” and “f_inv200” as significant prognostic factors (Supplementary Table 5). In multivariate analysis using features derived from beyond invasive margins, “f_im200” remained as an independent significant factor (Supplementary Table 6A). In case of features from inner tumor area, “f_wt” and “f_inv200” were identified as prognostic factors. (Supplementary Table 6B). Including all features, “f_wt” and “f_im200” were remained as statistically significant prognostic factors (Supplementary Table 6C). Thus, these two variables were included in further analysis adjusting with clinicopathologic variables.

In the Yonsei cohort, univariate analysis revealed that LVI (*p*=.009) and “f_im200” (*p*=.002) were significantly associated with PFS. In the multivariate analysis, “f_im200” was proven as an independent significant prognostic factor (HR 0.004 95% CI 0.0001–0.1502, *p*=.0028) (Table 2).

**Table 2.**
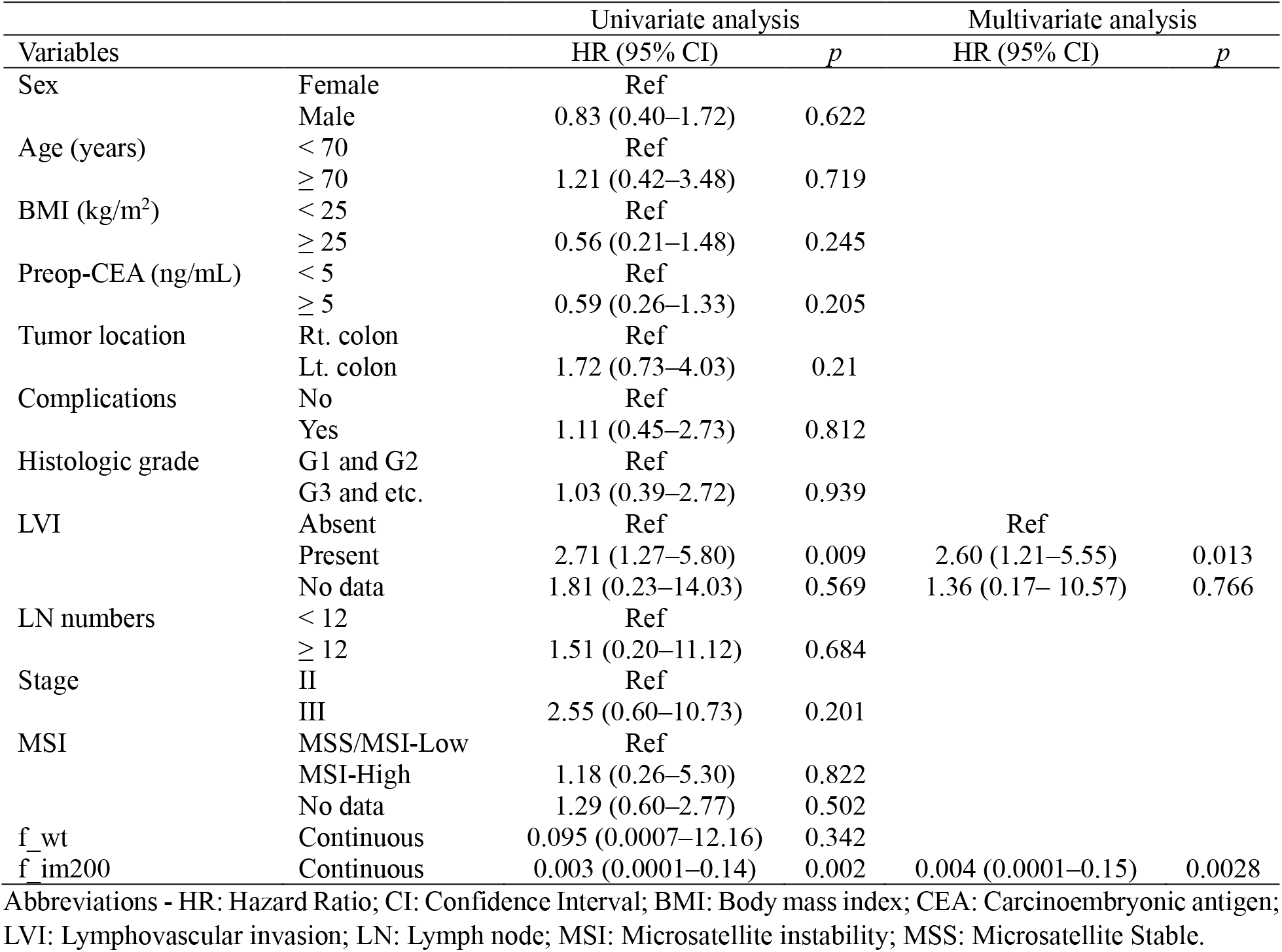
Univariate and multivariate analysis associated with progression-free survival in the Yonsei dataset (n=180)

### Validation of automated TILs features in the TCGA dataset

The median TILs density of spatial distribution of “f_wt” and “f_im200” were 0.0808 and 0.1085 respectively (Supplementary Table 7). Among the 268 patients from the included TCGA dataset, age, stage and “f_wt” and “f_im200” were proven to be significant prognostic factors in the univariate analysis. Factors *p* < 0.1 were entered into the multivariate analysis. Multivariate analysis revealed that “f_im200” retained prognostic significance for PFS (HR 0.031, [95% CI 0.001–0.645], *p*=.024) along with age (HR 1.956 [95% CI, 1.201–3.185], *p*=.006) and stage (I&II vs. IV, HR 3.194 [95% CI, 1.722–5.925], *p*=.002) (Table 3).

**Table 3.**
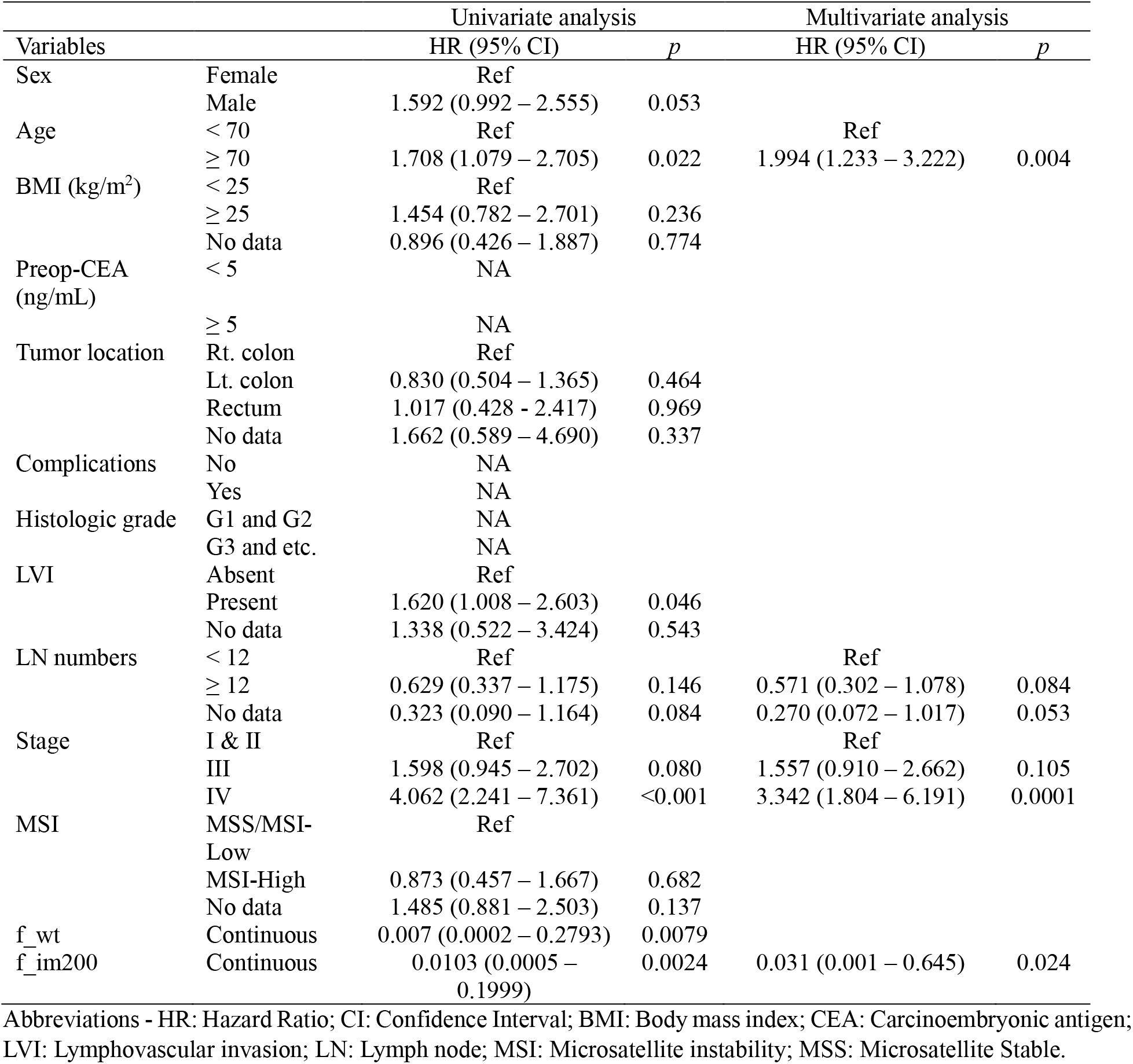
Univariate and multivariable analysis of factors associated with progression-free survival in the TCGA dataset (n=268)

### Interobserver agreement of KM grading for Yonsei and TCGA dataset by two expert pathologists

KM grading was manually performed by two pathologists for the Yonsei (n=180) and TCGA (n=249) dataset. Note that patients (n=19) from TCGA dataset with poor slide quality and/or slides which were difficult to manually find the invasive margins by pathologists were excluded for the analysis. The agreement of KM grading for each dataset by two pathologists was evaluated using kappa statistics.

Interobserver agreement of each KM grading was proven to be insignificant agreement in the Yonsei dataset (κ = .109) and in the TCGA dataset (κ = .121). When we repeated kappa statistics using KM-low (KM grading 0 and 1) versus KM-high (KM grading 2 and 3) groups, insignificant agreement was still observed in the Yonsei dataset (κ = .151), while, median agreement was observed in the TCGA dataset (κ = .404) (Supplementary Table 8).

### Comparison of patient subgrouping based on the DeepTILs and KM grading by the pathologists

We first investigated whether there are statistically significant differences of “f_im200” values across KM grading patient subgroups. The median values of “f_im200” according to the KM grading 0, 1, 2, and 3 by the pathologist 1 were 0.075, 0.142, 0.230 and 0.329 respectively in the Yonsei dataset (*p*<.001). The median values of “f_im200” according to the KM grading 1, 2, and 3 by the pathologist 2 were 0.055, 0.194 and 0.234 respectively in the Yonsei dataset (*p*<.001). There was no KM grading 0 by the pathologist 2 in the Yonsei dataset.

The median values of “f_im200” according to KM grading 0, 1, 2, and 3 by the pathologist 1 were 0.058, 0.109, 0.142 and 0.225 respectively in the TCGA dataset (*p*<.001) The median values of “f_im200” according to KM grading 0, 1, 2, and 3 by the pathologist 2 were 0.049, 0.077, 0.123 and 0.179 respectively in the TCGA dataset (*p*<.001) (Supplementary Figure 6). These results indicate higher KM grading patient subgroups carrying higher TILs densities of 200 μm IM layer.

We used the X-tile program to find an optimal cut-off value of “f_im200” in the Yonsei dataset and identified 0.14 as the cut-off value producing the largest χ2 in the Mantel–Cox test (Supplementary Figure 7). Based on this cut-off value, we divided patients from the Yonsei dataset into two subgroups (i.e., patients having higher than 0.14 of “f_im200” as a TILs high subgroup, otherwise low subgroup, respectively). The same cut-off value (0.14) was applied to identify TILs high and low subgroups for patients from the TCGA dataset.

Since 249 patients were graded by both pathologists, those 249 patients only included for the survival analysis. When we re-analyzed the multivariate analysis using the 249 selected patients in the TCGA dataset, f_im200 was again identified as an independent prognostic factor for PFS (HR 0.005, [95% CI 0.001–0.173], *p*=.003) (Supplementary Table 9). When we evaluated the association between the binary classification of DeepTILs and pathologists’ KM grading systems in these 249 patients, as the KM grading increased, the rate of high grading by DeepTILs also increased. (Supplementary Figure 8, 9)

### Prognostic utility of spatial TILs features based on DeepTILs and KM grading

We performed a Kaplan-Meier survival analysis, and found that patients of TILs high subgroup (e.g., patients with higher densities defined by the DeepTILs) showed better PFS compared with those patients belonged to the TILs low subgroup (A log rank test *p*=.0018) (Figure 2A). Interestingly, there were no survival differences between KM-low and KM-high groups either by the pathologist 1 (*p*=. 16) and the pathologist 2 (*p*=.66) respectively (Figure 2B and 2C).

**Figure 2.**
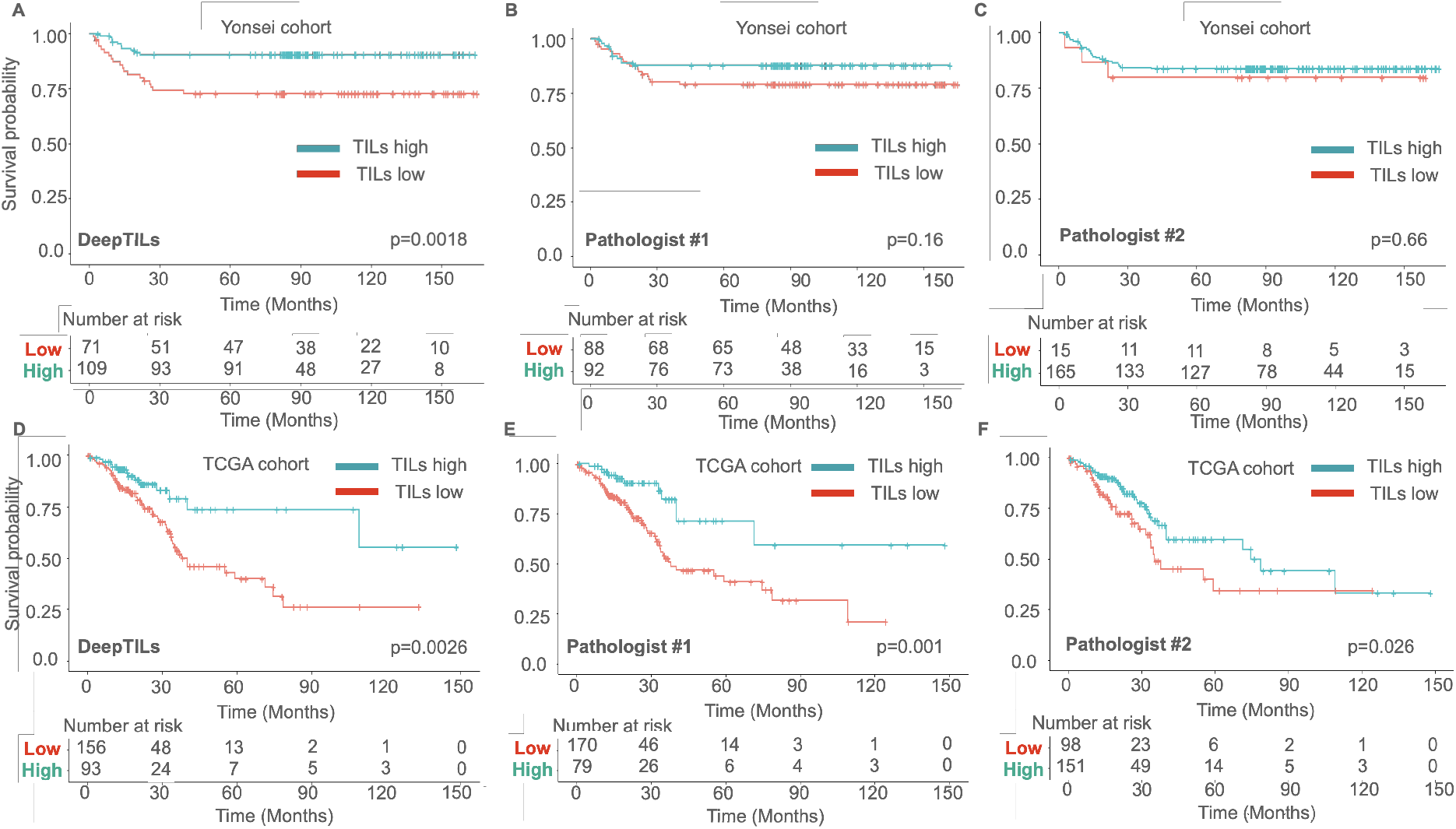
Kaplan Meier plots for patient subgroup analysis identified by DeepTILs and KM gradings by the pathologists. (A) TILs high and low subgroups identified by DeepTILs in Yonsei cohort (B-C) TILs high and low subgroups identified by KM gradings from the pathologists in Yonsei cohort. (D) TILs high and low subgroups identified by DeepTILs in TCGA cohort (E-F) TILs high and low subgroups identified by KM gradings from the pathologists in TCGA cohort.

A Kaplan-Meier survival analysis shows that those patients with higher densities defined by the DeepTILs showed better PFS when compared with those patients with lower densities (A log rank test *p*=.0026) (Figure 2D). The KM high group defined by the pathologist 1 and the pathologist 2 showed better PFS than the KM low groups (A log rank test *p*=.001 by the pathologist 1 and *p*=.026 by the pathologist 2) (Figure 2E and 2F).

### Combination of DeepTILs and pathologic grading and its prognostic effect

Lastly, we investigated whether integrating KM grading by the pathologists with TILs subgroups by DeepTILs could improve patient prognostication. Specifically, we group patients into four subgroups, TILs high and high, TILs high and low, TILs low and high, and TILs low and low by DeepTILs and the pathologists, respectively. We generate Kaplan-Meier plots for each patient’s KM grading with DeepTILs based subgroups and performed univariate and multivariate analysis of subgroups (Figure 3, Supplementary Table 10). Interestingly, patients belonged to TILs high subgroup by both KM grading and DeepTILs showed better PFS across datasets, while patients belonged to TILs low subgroup by both approaches showed poorest PFS. For example, on multivariate analysis using the TCGA dataset, we found that TILs high subgroups identified by both approaches show statistically significant PFS compared to TILs low subgroups identified by two approaches (e.g., TILs high subgroup by the pathologist #1 and DeepTILs: HR 0.372 [95% CI, 0.154-0.896], *p*=.027, and TILs high subgroup by the pathologist #2 and DeepTILs: HR 0.472 [95% CI, 0.230-0.968], *p*=.040, respectively) (Supplementary Table 10C and 10D). Another interesting observation is that patients assigned TILs low by KM grading but TILs high by DeepTILs showed a trend towards better PFS compared to TILs low subgroups by both approaches. However, patients assigned with TILs high by KM grading but TILs low by DeepTILs showed poorer PFS compared to those of TILs high subgroup by both approaches. This might indicate that incorporating TILs subgroups derived by DeepTILs with the pathologists’ KM grading could improve patient stratification compared to the KM grading alone.

**Figure 3.**
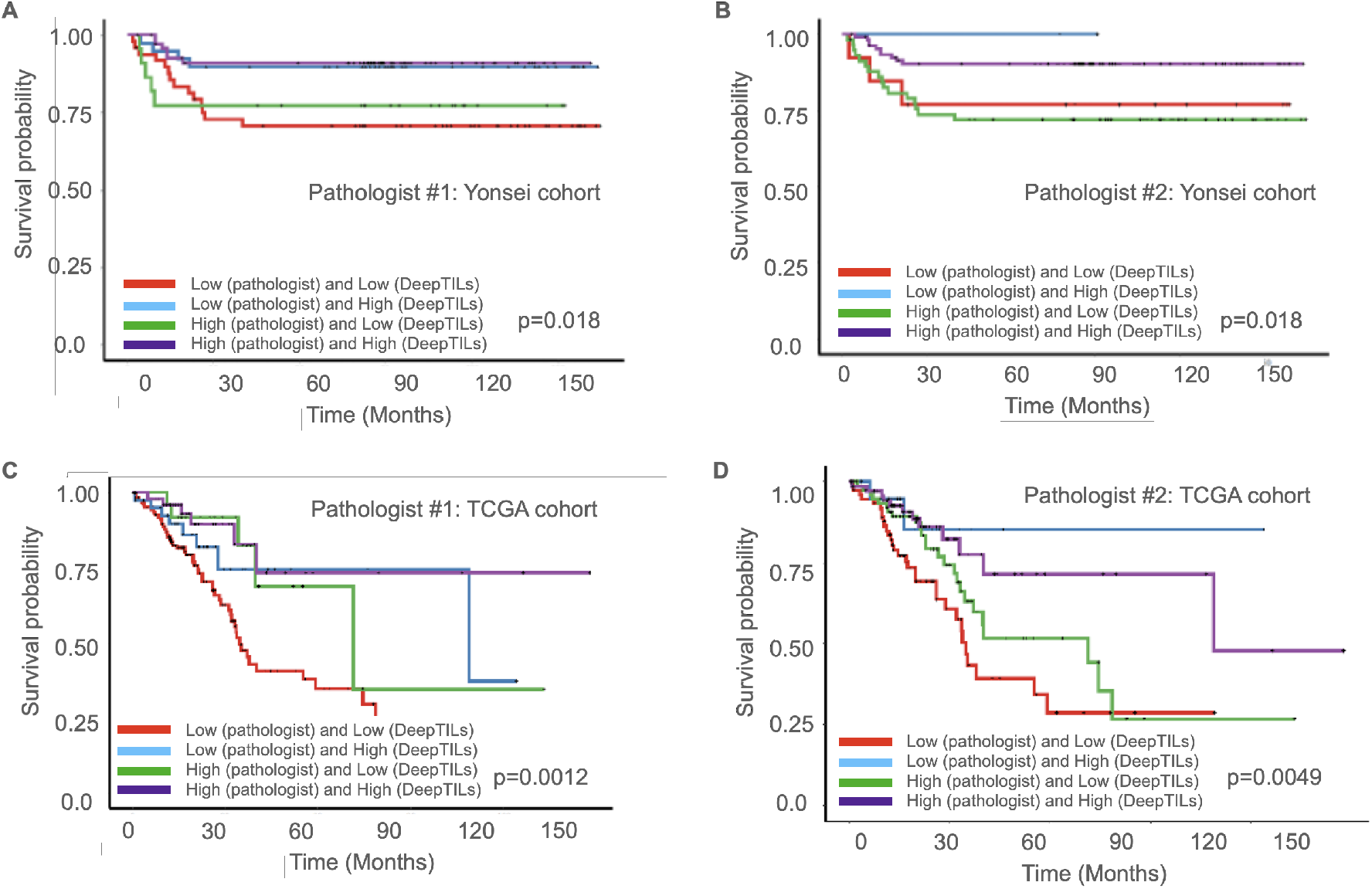
Kaplan-Meier survival analysis based on combination of subgroups derived from KM grading and DeepTILs. We assigned patients into four subgroups, TILs high and high, TILs high and low, TILs low and high, and TILs low and low by KM grading from two pathologists and DeepTILs in Yonsei and TCGA CRC cohorts, respectively. (A) Combination of patient subgroups by the pathologist #1 and DeepTILs in Yonsei cohort. (A) Combination of patient subgroups by the pathologist #2 and DeepTILs in Yonsei cohort. (A) Combination of patient subgroups by the pathologist #1 and DeepTILs in TCGA cohort. (A) Combination of patient subgroups by the pathologist #2 and DeepTILs in TCGA cohort.

## Discussion

We have shown that the DL models based on H&E-stained WSIs can quantify TILs densities at the invasive margins and overall regions of surgically resected CRC. In experiments using datasets from the Yonsei and TCGA cohorts, we showed that TILs densities quantified by DeepTILs could be used to identify subgroups of patients with CRC with distinct PFS difference. In particular, we performed comprehensive evaluation of TILs densities at various IMs as well as tumor core and whole tumor regions. Our subgroup analysis based on the automatic quantification of TILs densities in 200 μm IM layer (e.g., “f_im200”) by DeepTILs showed that the patient subgroup with high TILs densities in 200 μm IM have statistically significant different PFS outcome from the Yonsei and TCGA cohorts. However, the subgroup analysis-based on pathologists’ independent TILs grading only showed statistically significant different PFS outcome from the TCGA cohort not the Yonsei cohort.

Pathologist’s manual TILs grading on H&E or IHC stained slides are involved with manual tumor boundary, tumor invasive margin, etc. steps which are time consuming and labor intensive. In addition, intra-rater reliability of manual TILs grading by pathologists could make difficult universally deploy TILs analysis as a routine clinical practice. For example, the agreement of KM scoring using 4 grading system between the pathologists was insignificant in both the Yonsei and TCGA datasets indicating intra-rater issues for manual grading. The agreement of KM scoring using 2 grading system (e.g., patients grouped into two categories such as KM low versus high subgroups) showed that kappa value increased in the TCGA dataset, but no change observed in the Yonsei dataset, which indicate both 2 and 4 manual KM grading systems contain intra-rater issue.

One of strength of our approaches is to reduce inter-rater reliability by evaluating densities of TILs across various regions of tumors and at the invasive margins with less biases compared to human grading. Specifically, the deep learning approaches measured density of TILs of whole tumor region and invasive margins without manually selecting certain regions (e.g., manually selected representative or subset of tumor regions and/or invasive margins) which can reduce the inter-rater variability to measure TILs densities. In addition, our approaches are not limited to measure TILs given a certain region but can measure the different densities of TILs throughout tumor center and invasive margins and thus provide comprehensive prognostic evaluation across whole regions of tumor core and boundary using H&E based WSIs.

The pathologists revisited cases from TCGA cohort which have different TILs high or low grading at tumor invasive margin by KM grading and DeepTILs, We found that the disagreement of subgroups was largely due to lack of consensus of tumor invasive margin definition (Supplementary Figure 10). For instance, most of cases with TILs high subgroup by the pathologists have high level TILs quantification within tumors by DeepTILs. However, those cases have few TILs present in tumor invasive margin defined by DeepTILs, thus graded as TILs low (Supplementary Figure 10C and 10D). Similarly, TILs low subgroup by KM grading have few TILs within tumors, while those few TILs were present in tumor invasive margin defined by DeepTILs (Supplementary Figure 10A and 10B). We also found that some inflammatory cells in necrotic debris and/or fibrosis tissue from few cases were detected as TILs by DeepTILs (Supplementary Figure 10A and 10B). While this false positive could lead the discrepancy of patient subgrouping by two approaches, the DL approaches could be further improved by learning these patterns.

Recent meta-analysis showed that manual TILs analysis of certain types of T-cells (e.g., CD4, CD8, etc.) in tumor center, stroma and at the IM based on IHC staining from multiple studies found prognostic information of those manual TILs densities associated with overall survival (OS) and disease-free survival.(2) The discrepancies of findings (i.e., no statistical significance of TILs densities in tumor center derived by DeepTILs) compared to recent meta-analysis could be due to that our approaches only utilized H&E stained WSIs which we cannot take into account densities of certain type of T-cells within tumor region and at invasive margins. Also our discovery cohort, the Yonsei cohort, is collected from patients who went to surgical resection from a single institute and could poise ethnic specific disease outcome and/or morphologic differences across ethnicities.(19, 20) Our approaches are general approaches which can utilize IHC stained WSIs to quantify densities of TILs of certain types of T-cells in tumor region, core and at invasive margins. We plan to generate and collect IHC stained WSIs with CD3, CD8 and FOXP3 to quantify TILs density and evaluate prognostic information of TILs features from a certain type T-cells across tumor center, core and at invasive margins. With additional larger training and testing datasets from multi center and international institutes, we will also further validate our findings whether TILs densities in 200 μm IM layer could serve as a robust prognostic biomarker for patients with CRC. Another limitation was we did not incorporate morphologic features present in tumor and stroma regions in addition to TILs densities at various tumor regions and IMs to correlate with patient’s outcome. There are several new studies showing that H&E stained WSIs-based deep learning models utilizing morphologic features from WSIs can accurately predict survival of patients with CRC. Incorporating such morphologic features with TILs densities could further improve patient’s prognostication. Lastly, while there are some agreements between automated and manual TILs grading from the deep learning approaches and pathologists, we did not attempt to incorporate manual TILs grading into the patient subgroup analysis or to use the deep learning model as an assistant for pathologists’ TILs grading. Recent study showed that incorporating the deep learning model to predict MSI status based on H&E stained images could provide labour and cost saving benefits.(21) Similarly, proper integration of the deep learning models to access TILs grading could potentially provide similar benefits.

Taken together, we developed H&W stained WSIs based deep learning approaches for TILs spatial detection and quantification. Our analyses indicate that the use of the DL approaches for TILs grading could be used to identify patient subgroups with distinct PFS. The analyses also indicated that the use of the DL approaches could be used to address the interobserver disagreements of H&E based TILs evaluation by pathologists. Finally, we showed that incorporating the DL based TILs subgroup with the KM grading could improve patient prognostication. Our results should warrant to further test and validate the deep learning approaches for spatial TILs quantification through a larger number and diverse datasets as well as comparing the IHC based scoring system to help to select CRC patients with high or low risk.

## Acknowledgements

The authors thank Medical Illustration & Design, part of the Medical Research Support Services of Yonsei University College of Medicine, for all artistic support related to this work.

## Funding

This work was supported by the National Research Foundation of Korea(NRF) grant funded by the Korea government(MSIT) (No. 2020R1G1A1102555).

## Conflicts of interest

THH is co-founder of KURE.AI and received consulting fees and research funding from AITRICS.

## Notes

https://github.com/hwanglab/TILs_Analysis

### Competing Interest Statement

THH received consulting fees from AITRICS. THH received research funding from AITRICS through the insititute. THH is co-founder of KURE.AI.

