## Supplementary Information for "Spatial analysis of tumor infiltrating lymphocytes based on deep learning using histopathology image to predict progression-free survival in colorectal cancer"

### Supplementary files

**Supplementary Table 1. Grid Search of parameter settings for Resnet18 based Tumor detectors**

| Parameters | Frozen Ratios | Optimizers | Learning Rates | Batch Sizes |
|---|---|---|---|---|
| Settings | {0%, 80%} | {‘sgd’,’adam’} | {‘0.001’,’0.0001’} | {4,16,64} |

: The frozen ratios refer to the percentages of early trainable layers that were frozen during transfer learning. We considered only early trainable convolution and feedforward layers, and did not count or freeze batch normalization or activation layers. By using different parameter settings, 24 different tumor detectors were trained

**Supplementary Table 2. Grid Search of parameter settings for finding the best tumor-infiltrating lymphocytes (TILs) detector**

| Parameters | Models | Frozen Ratios | Optimizers | Learning Rates | Batch Sizes |
|---|---|---|---|---|---|
| Settings | {‘resnet18’,’resnet34’, ‘shufflenet’} | {0%, 80%} | {‘sgd’,’adam’} | {‘0.001’,’0.0001’, ‘0.00001’} | {4,16,32,64} |

: By using different parameter settings, 144 different TILs detectors were trained.

**Supplementary Table 3. Descriptions of computed tumor-infiltrating lymphocytes (TILs) spatial distribution variables**

| Invasive margins | Descriptions | Tumor regions | Descriptions |
|---|---|---|---|
| f_im200 | TILs densities at 200um invasive margin layer | f_wt | TILs densities at whole tumor region |
| f_im300 | TILs densities at 300um invasive margin layer | f_inv200 | TILs densities at 200um inverse invasive margin layer |
| f_im400 | TILs densities at 400um invasive margin layer | f_tc | TILs densities at tumor core region |
| f_im500 | TILs densities at 500um invasive margin layer | f_tc2 | TILs densities at tumor regions excluding the area of inverse 200 invasive margin layer |

**Supplementary Table 4. Composition of tumor-infiltrating lymphocytes (TILs) according to spatial distribution in the Yonsei dataset**

| | Min | 25 percentile | Median | 75 percentile | Max | Mean |
|---|---|---|---|---|---|---|
| f_wt | 0.0116 | 0.0462 | 0.0857 | 0.1335 | 0.6814 | 0.1109 |
| f_im200 | 0 | 0.1073 | 0.2002 | 0.3003 | 0.8850 | 0.2206 |
| f_im300 | 0 | 0.1009 | 0.1870 | 0.2855 | 0.8790 | 0.2120 |
| f_im400 | 0 | 0.0967 | 0.1761 | 0.2715 | 0.8630 | 0.2037 |
| f_im500 | 0 | 0.0991 | 0.1684 | 0.2629 | 0.8359 | 0.1975 |
| f_inv200 | 0.0218 | 0.0965 | 0.1627 | 0.2426 | 0.7482 | 0.1795 |
| f_tc2 | 0.0023 | 0.0341 | 0.0616 | 0.1174 | 0.6694 | 0.093 |
| f_tc | 0 | 0.0206 | 0.0447 | 0.0993 | 0.6623 | 0.0847 |

**Supplementary Table 5. Univariate Cox proportional hazard ratio of progression-free survival in the Yonsei dataset**

| | Beta | HR (95% CI) | Wald test | *p* |
|---|---|---|---|---|
| f_wt | -2.35 | 0.0953 (0.0007 – 12.16) | 0.9 | 0.342 |
| f_im200 | -5.5518 | 0.0038 (0.0001 – 0.1475) | 8.95 | 0.0027 |
| f_im300 | -5.6362 | 0.0035 (8.15e-05 – 0.1561) | 8.55 | 0.0034 |
| f_im400 | -5.9216 | 0.0026 (5.034e-05 – 0.1427) | 8.53 | 0.0035 |
| f_im500 | - 6.1501 | 0.0021 (3.405e-05 – 0.1336) | 8.49 | 0.0035 |
| f_inv200 | -5.3144 | 0.0049 (6.643e-05 – 0.3644) | 5.85 | 0.0155 |
| f_tc2 | -1.3655 | 0.2553 (0.0028 – 22.95) | 0.35 | 0.552 |
| f_tc | -1.2733 | 0.2799 (0.0043 – 17.86) | 0.36 | 0.548 |

**Supplementary Table 6. Multivariate analysis of factors associated with progression-free survival using spatial tumor-infiltrating lymphocytes (TILs) densities in the Yonsei dataset**

6A) Multivariate analysis including features derived from beyond invasive margins

| | HR (95% CI) | *p* |
|---|---|---|
| f_im200 | 0.00388 (0.0001021 – 0.1475) | 0.00278 |
| f_im300 | | |
| f_im400 | | |
| f_im500 | | |

6B) Multivariate analysis including features derived from inner tumor area

| | HR (95% CI) | *p* |
|---|---|---|
| f_wt | 1.009e+13 (3.439e+03 – 2.962e+22) | 0.007101 |
| f_inv200 | 5.743e-09 (1.446e-13 – 2.281e-04) | 0.000445 |
| f_tc2 | | |
| f_tc | 1.343e-06 (1.638e-12 – 1.102e+00) | 0.051656 |

6C) Multivariate analysis including features derived from beyond invasive margins and inner tumor area

| | HR (95% CI) | *p* |
|---|---|---|
| f_wt | 6.615e+02 (1.961e+00 – 2.231e+05) | 0.028760 |
| f_im200 | 1.720e-04 (1.239e-06 – 2.389e-02) | 0.000574 |
| f_im300 | | |
| f_im400 | | |
| f_im500 | | |
| f_inv200 | | |
| f_tc2 | | |
| f_tc | | |

**Supplementary Table 7. Composition of tumor-infiltrating lymphocytes (TILs) according to spatial distribution in the TCGA dataset**

| | Min | 25 percentile | Median | 75 percentile | Max | Mean |
|---|---|---|---|---|---|---|
| f_wt | 0 | 0.0373 | 0.0808 | 0.1364 | 0.9614 | 0.1059 |
| f_im200 | 0 | 0.0548 | 0.1085 | 0.1885 | 0.6936 | 0.1885 |

**Supplementary Table 8. Comparison of Kappa value between the two pathologists using validated slides in the Yonsei dataset (n=180) and the TCGA dataset (n=249)**

| | KM grading 0, 1, 2, and 3 | | KM grading low vs. high | |
|---|---|---|---|---|
| | κ | *p* | κ | *p* |
| Yonsei dataset | 0.109 | 0.00625 | 0.151 | 0.000322 |
| TCGA dataset | 0.121 | 0.000106 | 0.404 | 4.31e-14 |

**Supplementary Table 9. Univariate and multivariable analysis of factors associated with PFS in the TCGA dataset after selection by the pathologists (n=249)**

| | | Univariate analysis | | Multivariate analysis | |
|---|---|---|---|---|---|
| Variables | | HR (95% CI) | *p* | HR (95% CI) | *p* |
| Sex | Female | Ref | | | |
| | Male | 1.616 (0.992–2.63) | 0.053 | | |
| Age | < 70 | Ref | | Ref | |
| | ≥ 70 | 1.63 (1.014–2.62) | **0.043** | 1.888 (1.148–3.103) | **0.012** |
| BMI (kg/m$^2$) | < 25 | Ref | | | |
| | ≥ 25 | 1.408 (0.757–2.621) | 0.280 | | |
| | No data | 0.746 (0.341–1.633) | 0.465 | | |
| Preop-CEA (ng/mL) | < 5 | NA | | | |
| | ≥ 5 | NA | | | |
| Tumor location | Rt. colon | Ref | | | |
| | Lt. colon | 0.921 (0.553–1.535) | 0.753 | | |
| | Rectum | 1.073 (0.447–2.568) | 0.874 | | |
| | No data | 1.918 (0.676–5.443) | 0.221 | | |
| Complications | No | NA | | | |
| | Yes | NA | | | |
| Histologic grade | G1 and G2 | NA | | | |
| | G3 and etc. | NA | | | |
| LVI | Absent | Ref | | | |
| | Present | 1.643 (1.008–2.679) | **0.046** | | |
| | No data | 1.372 (0.534–3.520) | 0.511 | | |
| LN numbers | < 12 | Ref | | Ref | |
| | ≥ 12 | 0.606 (0.323–1.136) | 0.118 | 0.568 (0.299–1.078) | 0.083 |
| | No data | 0.324 (0.090–1.164) | 0.084 | 0.281 (0.074–1.058) | 0.060 |
| Stage | I & II | Ref | | Ref | |
| | III | 1.622 (0.940–2.797) | 0.082 | 1.457 (0.833–2.550) | 0.186 |
| | IV | 4.098 (2.241–7.493) | **<0.001** | 3.053 (1.628–5.726) | **0.0005** |
| MSI | MSS/MSI-Low | Ref | | | |
| | MSI-High | 0.777 (0.387–1.559) | 0.478 | | |
| | No data | 1.540 (0.904–2.624) | 0.112 | | |
| f_wt | Continuous | 0.005 (0.0001–0.2485) | **0.0074** | | |
| f_im200 | Continuous | 0.0017 (6.139e-05–0.050) | **0.0002** | 0.005 (0.0001–0.173) | **0.003** |

Abbreviations - HR: Hazard Ratio; CI: Confidence Interval; BMI: Body mass index; CEA: Carcinoembryonic antigen; LVI: Lymphovascular invasion; LN: Lymph node; MSI: Microsatellite instability; MSS: Microsatellite Stable.

NA: Not available

**Supplementary Table 10. Multivariate analysis of factors associated with PFS in the Yonsei dataset and TCGA dataset using combination of KM grading by the pathologists and DeepTILs**

**10A) Combination of Pathologist 1 and DeepTILs in the Yonsei dataset**

| | | Univariate analysis | | Multivariate analysis | |
|---|---|---|---|---|---|
| Variables | | HR (95% CI) | *p* | HR (95% CI) | *p* |
| LVI | Absent | Ref | | Ref | |
| | Present | 2.719 (1.273–5.807) | **0.009** | 2.838 (1.324–6.084) | **0.007** |
| | No data | 1.812 (0.234–14.03) | 0.569 | 1.374 (0.173–10.866) | 0.763 |
| Combination of manual grading and DeepTILs | C1 | Ref | | Ref | |
| | C2 | 0.327 (0.107–0.993) | **0.048** | 0.288 (0.094–0.881) | **0.029** |
| | C3 | 0.833 (0.300–2.315) | 0.727 | 0.869 (0.310–2.433) | 0.789 |
| | C4 | 0.282 (0.108–0.735) | **0.009** | 0.289 (0.110–0.758) | **0.011** |

C1: Low grading by the pathologist and low grading by the DeepTILs, C2: Low grading by the pathologist and high grading by the DeepTILs, C3: High grading by the pathologist and low grading by the DeepTILs, C4: High grading by the pathologist and high grading by the DeepTILs

**10B) Combination of Pathologist 2 and DeepTILs in the Yonsei dataset**

| | | Univariate analysis | | Multivariate analysis | |
|---|---|---|---|---|---|
| Variables | | HR (95% CI) | *p* | HR (95% CI) | *p* |
| LVI | Absent | Ref | | Ref | |
| | Present | 2.719 (1.273–5.807) | **0.009** | 2.719 (1.273–5.807) | **0.009** |
| | No data | 1.812 (0.234–14.03) | 0.569 | 1.812 (0.234–14.03) | 0.569 |
| Combination of manual grading and DeepTILs | C1 | Ref | | | |
| | C2 | 6.217e-08 (0–Inf) | 0.998 | | |
| | C3 | 1.201e+00 (0.35–4.123) | 0.771 | | |
| | C4 | 3.736e-01 (0.102–1.358) | 0.135 | | |

C1: Low grading by the pathologist and low grading by the DeepTILs, C2: Low grading by the pathologist and high grading by the DeepTILs, C3: High grading by the pathologist and low grading by the DeepTILs, C4: High grading by the pathologist and high grading by the DeepTILs

**10C) Combination of Pathologist 1 and DeepTILs in the TCGA dataset (n=249)**

| | | Univariate analysis | | Multivariate analysis | |
|---|---|---|---|---|---|
| Variables | | HR (95% CI) | *p* | HR (95% CI) | *p* |
| Age | < 70 | Ref | | Ref | |
| | ≥ 70 | 1.63 (1.014–2.62) | **0.043** | 1.884 (1.147–3.096) | **0.012** |
| LVI | Absent | Ref | | | |
| | Present | 1.643 (1.008–2.679) | **0.046** | | |
| | No data | 1.372 (0.534–3.520) | 0.511 | | |
| LN numbers | < 12 | Ref | | Ref | |
| | ≥ 12 | 0.606 (0.323–1.136) | 0.118 | 0.555 (0.293–1.052) | 0.071 |
| | No data | 0.324 (0.090–1.164) | 0.084 | 0.264 (0.069–1.006) | 0.051 |
| Stage | I & II | Ref | | Ref | |
| | III | 1.622 (0.940–2.797) | 0.082 | 1.478 (0.834–2.618) | 0.180 |
| | IV | 4.098 (2.241–7.493) | **<0.001** | 3.089 (1.617–5.900) | **0.0006** |
| Combination of manual grading and DeepTILs | C1 | Ref | | Ref | |

| | | | | | |
|---|---|---|---|---|---|
| | C2 | 0.480 (0.227–1.016) | 0.055 | 0.686 (0.312–1.505) | 0.146 |
| | C3 | 0.370 (0.147–0.929) | **0.034** | 0.411 (0.162–1.038) | 0.059 |
| | C4 | 0.271 (0.116–0.634) | **0.002** | 0.372 (0.154–0.896) | **0.027** |

C1: Low grading by the pathologist and low grading by the DeepTILs, C2: Low grading by the pathologist and high grading by the DeepTILs, C3: High grading by the pathologist and low grading by the DeepTILs, C4: High grading by the pathologist and high grading by the DeepTILs

**10D) Combination of Pathologist 2 and DeepTILs in the TCGA dataset (n=249)**

| | | Univariate analysis | | Multivariate analysis | |
|---|---|---|---|---|---|
| Variables | | HR (95% CI) | *p* | HR (95% CI) | *p* |
| Age | < 70 | Ref | | Ref | |
| | ≥ 70 | 1.63 (1.014–2.62) | **0.043** | 1.945 (1.181–3.203) | **0.008** |
| LVI | Absent | Ref | | | |
| | Present | 1.643 (1.008–2.679) | **0.046** | | |
| | No data | 1.372 (0.534–3.520) | 0.511 | | |
| LN numbers | < 12 | Ref | | Ref | |
| | ≥ 12 | 0.606 (0.323–1.136) | 0.118 | 0.551 (0.291–1.042) | 0.066 |
| | No data | 0.324 (0.090–1.164) | 0.084 | 0.292 (0.076–1.119) | 0.072 |
| Stage | I & II | Ref | | Ref | |
| | III | 1.622 (0.940–2.797) | 0.082 | 1.388 (0.781–2.468) | 0.263 |
| | IV | 4.098 (2.241–7.493) | **<0.001** | 3.136 (1.635–6.015) | **0.0005** |
| Combination of manual grading and DeepTILs | C1 | Ref | | Ref | |
| | C2 | 0.253 (0.060–1.064) | 0.060 | 0.346 (0.076–1.511) | 0.158 |
| | C3 | 0.632 (0.372–1.072) | 0.088 | 0.636 (0.371–1.093) | 0.101 |
| | C4 | 0.348 (0.178–0.680) | **0.002** | 0.472 (0.230–0.968) | **0.040** |

C1: Low grading by the pathologist and low grading by the DeepTILs, C2: Low grading by the pathologist and high grading by the DeepTILs, C3: High grading by the pathologist and low grading by the DeepTILs, C4: High grading by the pathologist and high grading by the DeepTILs

A)

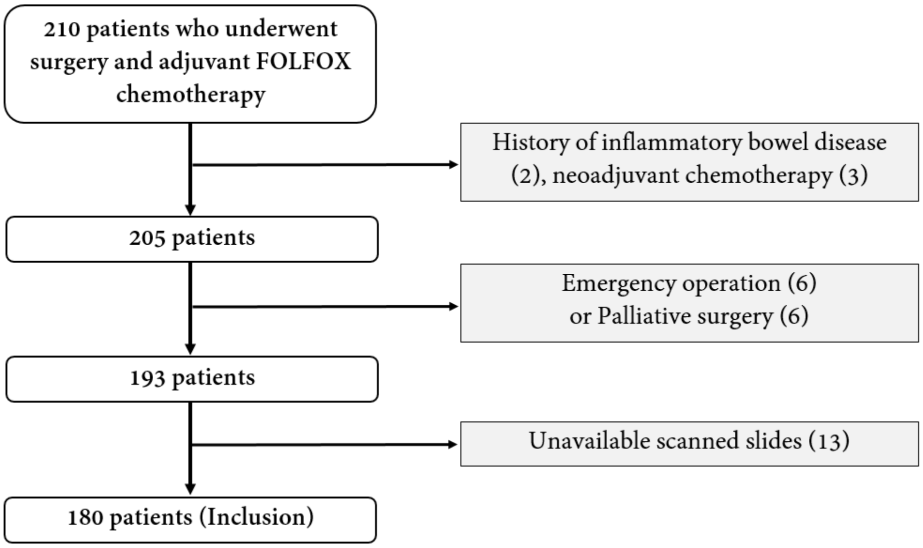


B)

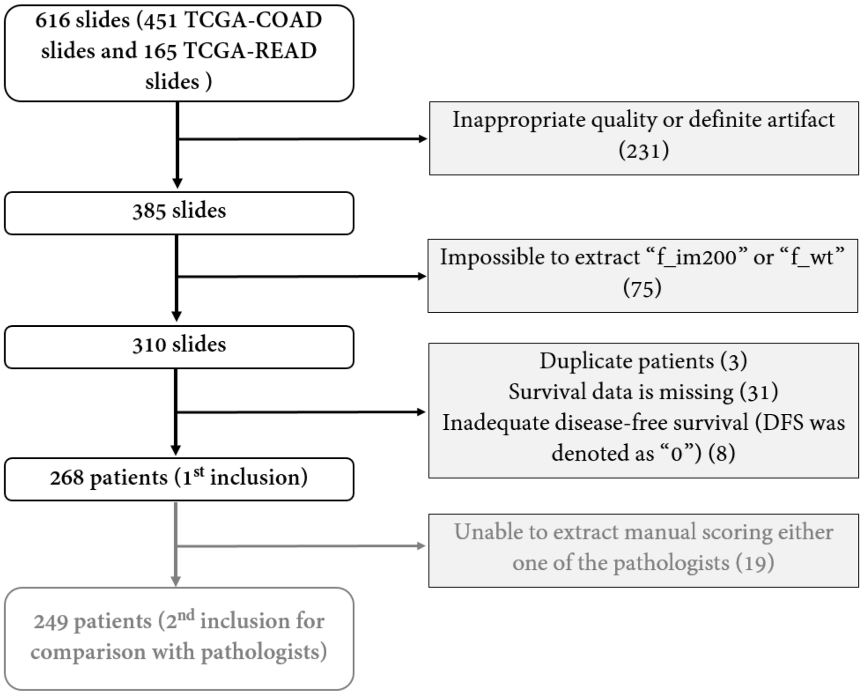


**Supplementary Figure 1. Patient inclusion of this study in the Yonsei dataset (A) and the TCGA dataset (B)**

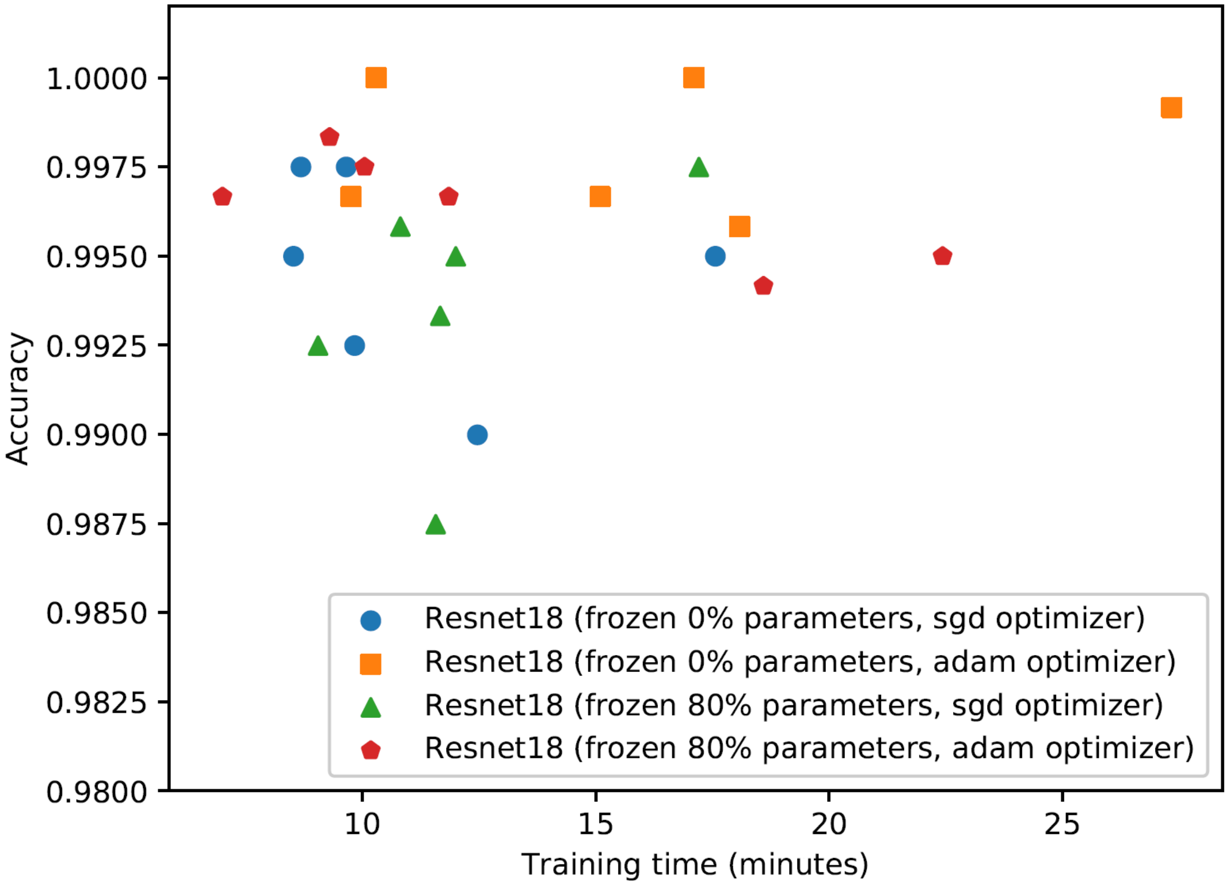


**Supplementary Figure 2. Independent testing on tumor detectors**

: Accuracies of 24 trained tumor detectors during the hyper-parameter search. In the figure, the horizontal axis corresponds to the training times of different models, while the vertical axis corresponds to testing accuracies of different models. Note that different symbols (i.e., circles, squares, triangles and pentagons) represent Restnet18 models with different parameter settings in terms of frozen ratios and optimizers. Each symbol (e.g., square) has 6 copies, which corresponds to different parameter settings in terms of batch sizes and learning rates. It could be found that two Resent18 models with 0% frozen ratios and adam optimizers provide the highest testing accuracies.

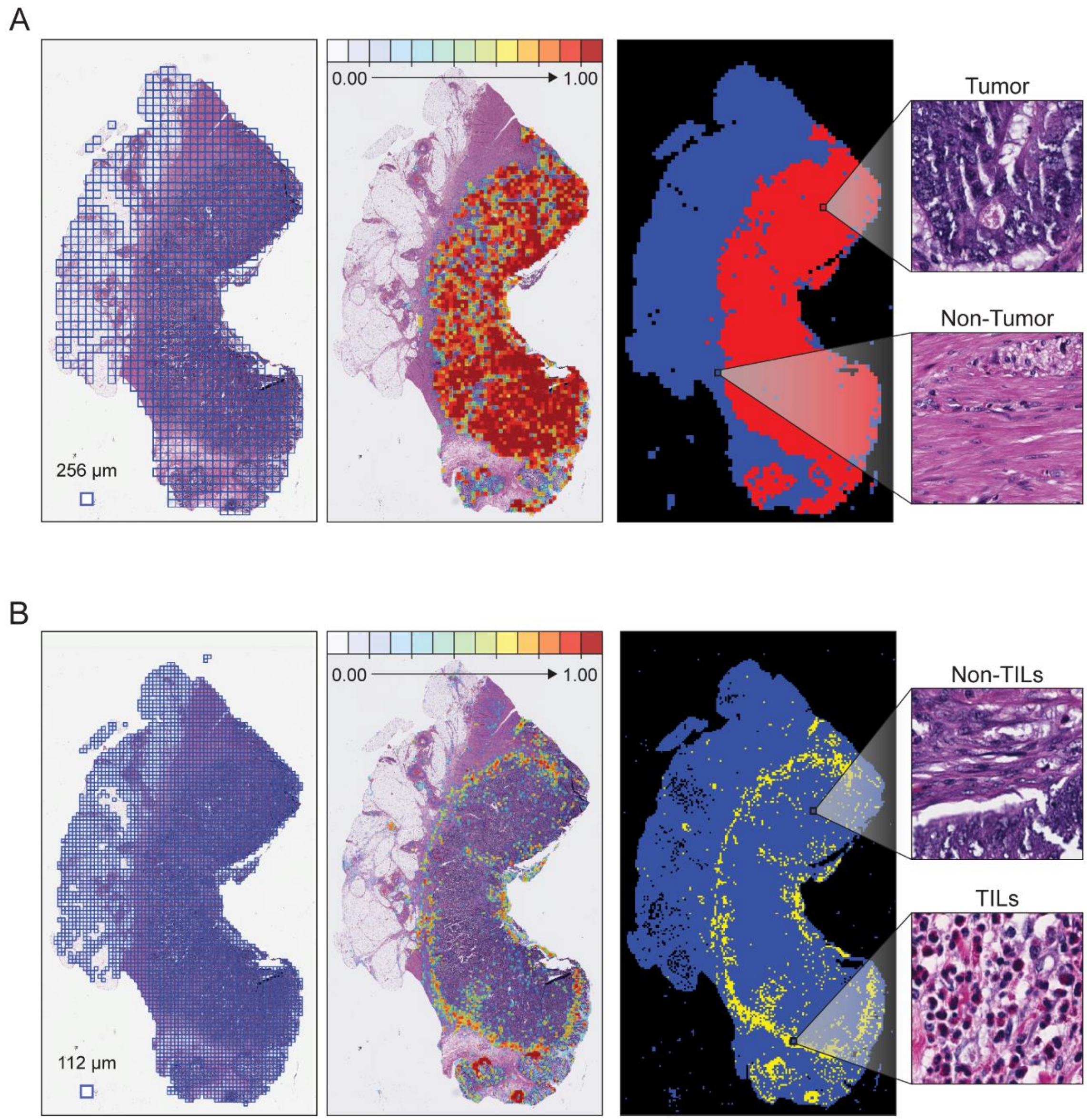


**Supplementary Figure 3. Heatmaps of predicted tumor cells and TILs.**

: (A) Tiled images are used to predict the probability of tumor cell by DeepTILs. Red color indicates higher probability of the tile carrying tumor cells. (B) Predicted TILs in WSIs. Red color indicates higher probability of tile carrying TILs.

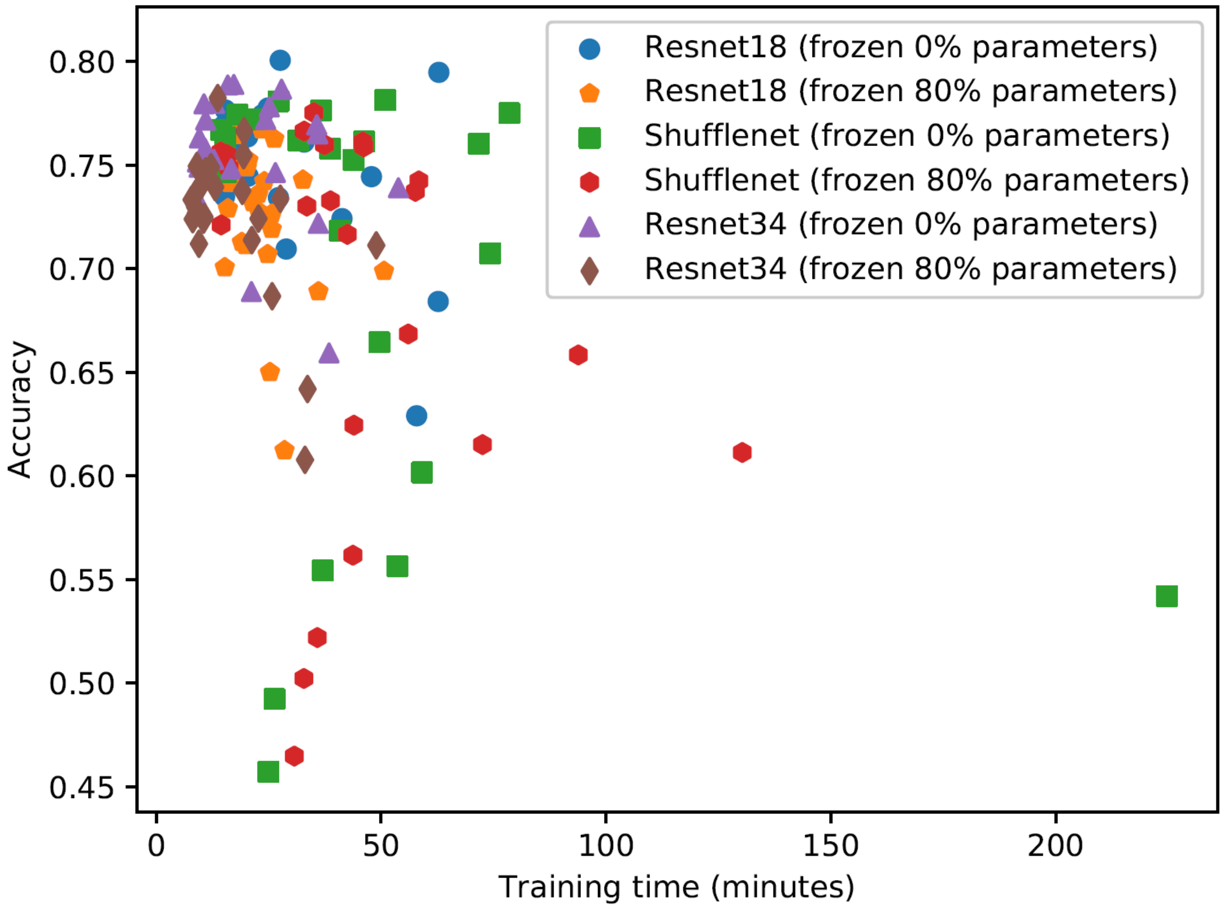


**Supplementary Figure 4. Independent testing on TILs detectors**

**:** Accuracies of 144 trained TILs detectors during the hyper-parameter search. In the figure, the horizontal axis corresponds to the training times of different models, while the vertical axis corresponds to testing accuracies of different models. Note that different symbols (e.g., circles, squares, triangles and pentagons) represent different models with different frozen ratios. Each symbol (e.g., square) has 24 copies, which corresponds to different parameter settings in terms of optimizers, batch sizes and learning rates. It could be found that two Resnet18 models with 0% frozen ratios provide noticeable higher accuracies than other models.

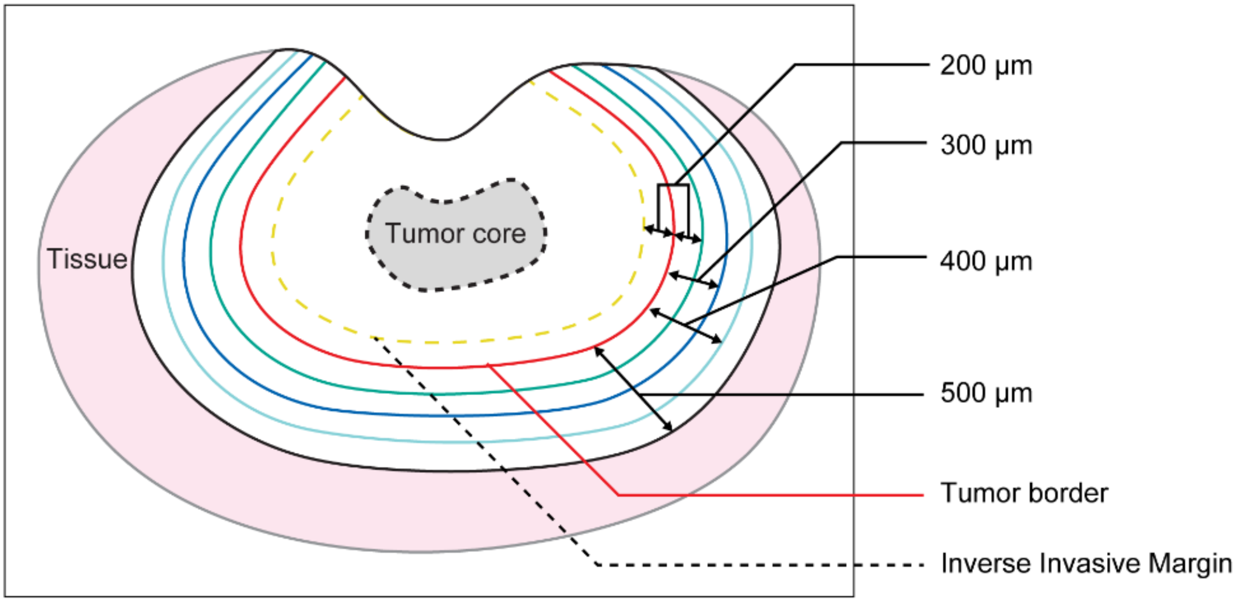


**Supplementary Figure 5.** Schematic region of interest selection based on tumor region

: We first define tumor border then select various tumor invasive margins and tumor core region. Tumor core is defined as a center region within a tumor. Tumor border region and various invasive margins were defined as in this diagram.

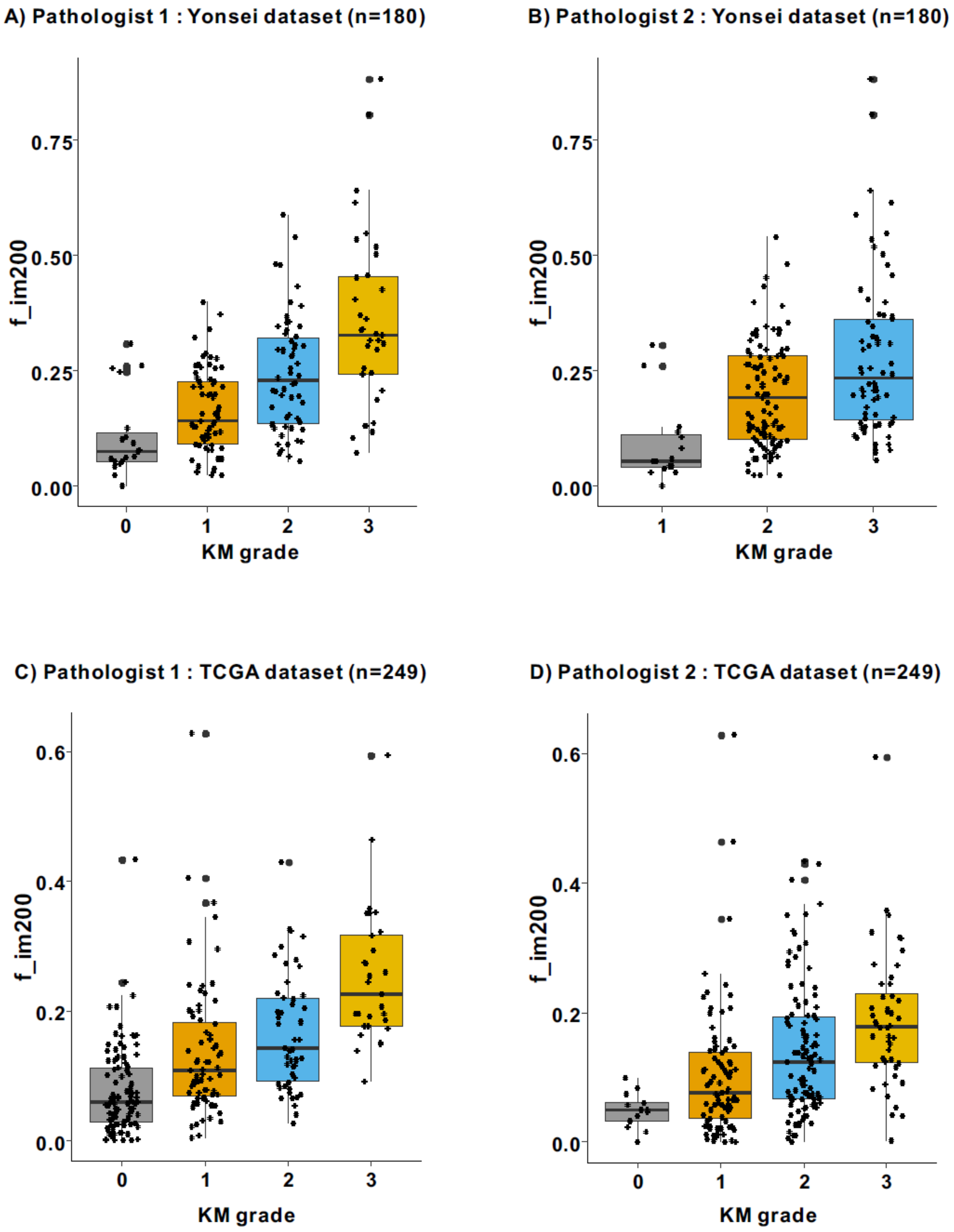


**Supplementary Figure 6. Distribution of f_im200 according to KM grading measured by the pathologist 1 and 2 using the Yonsei and the TCGA dataset**

A) Association diagram B) Cut-off value of the “f_im200”

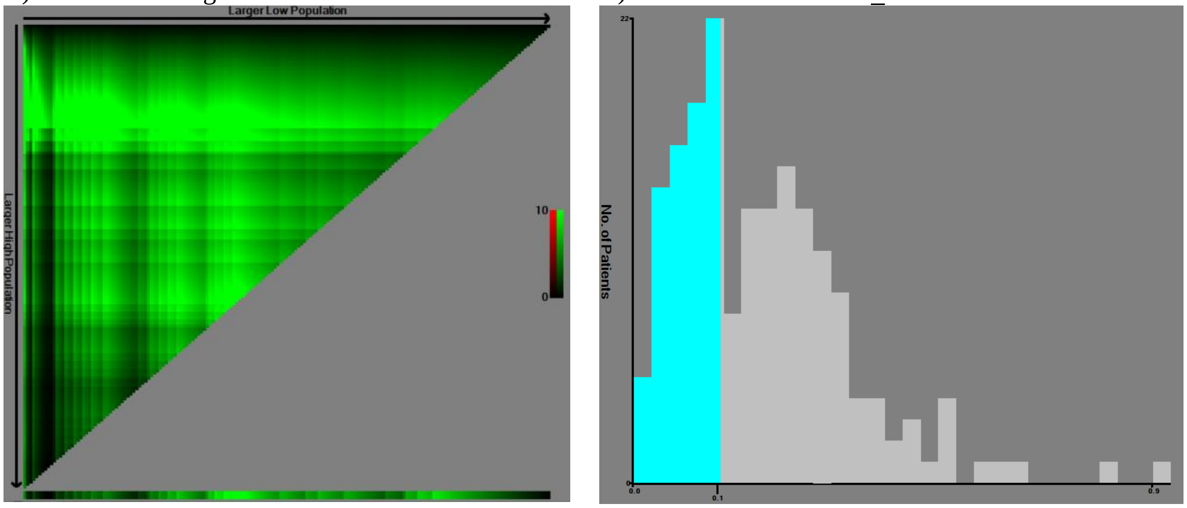


**Supplementary Figure 7. Determining cut-off values of “f_im200” using X-tile program in the Yonsei dataset (n=180)**

**:** X-tile plots of the “f_im200” and the points of the variable coloration of the plot represents the strength of the association at each division ranging from low (dark, black) to high (bright, red or green). Green represents a direct association between the expression levels and survival of the variables, whereas red represents an inverse association.(1)

The optimal cut-off value (0.14) was defined as the values that produced the largest $\chi 2$ in the Mantel–Cox test. Patients were divided into the high and low DeepTILs subgroups based on this value on the following Kaplan-Meier survival analysis in the Yonsei and TCGA dataset respectively.

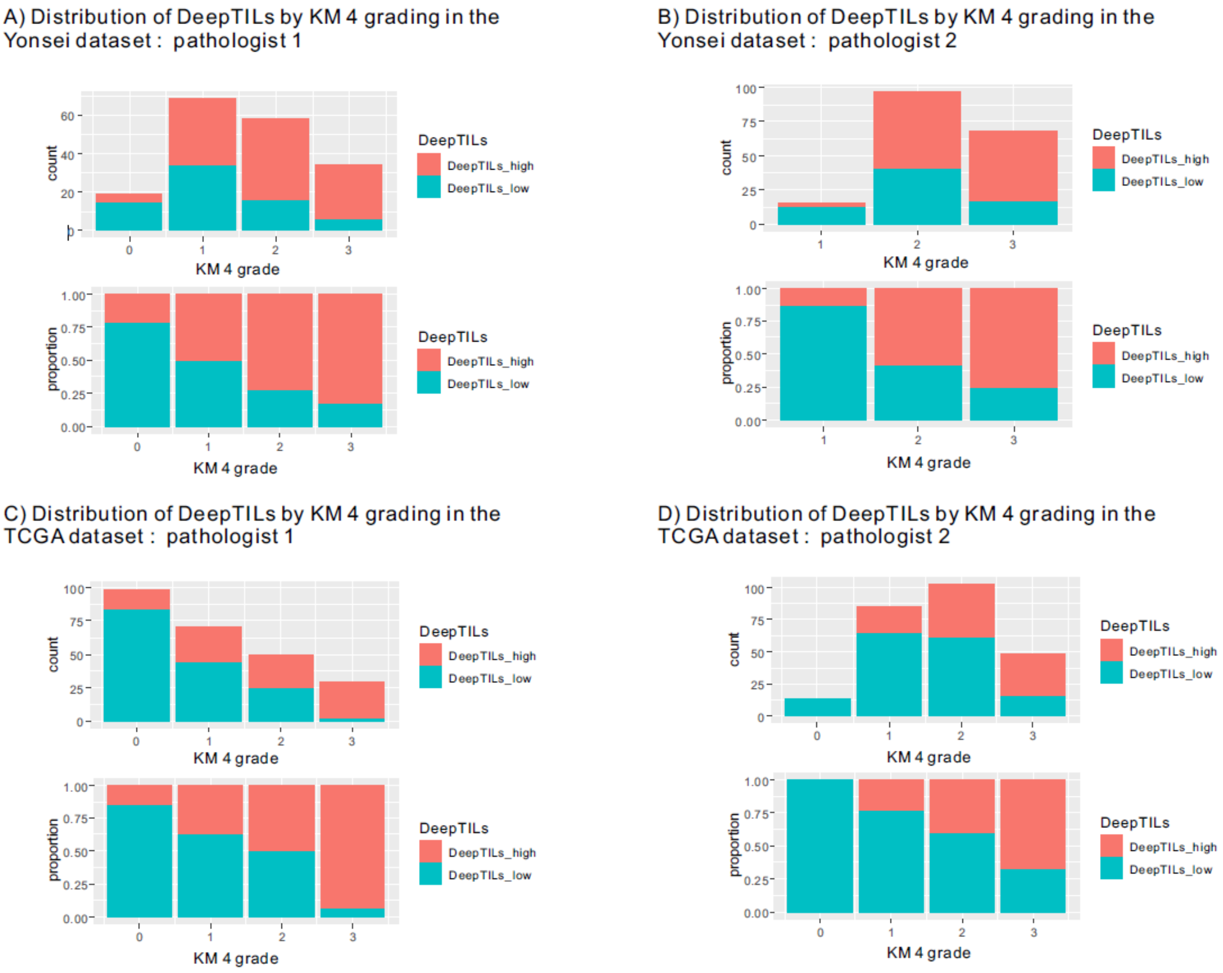


**Supplementary Figure 8. Distribution of DeepTILs by KM 4 grading by the two pathologists**

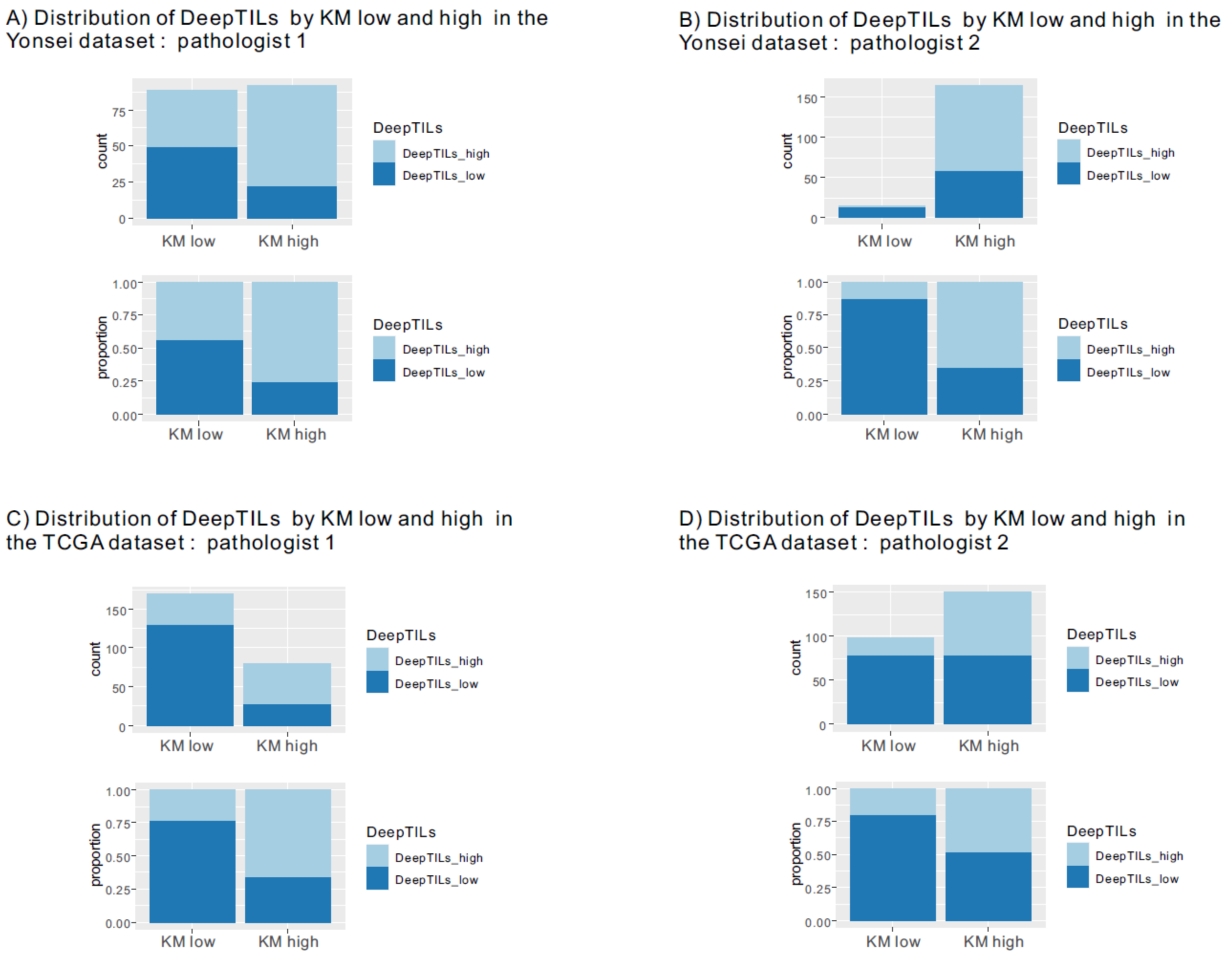


**Supplementary Figure 9. Distribution of DeepTILs by KM low and high by the two pathologists**

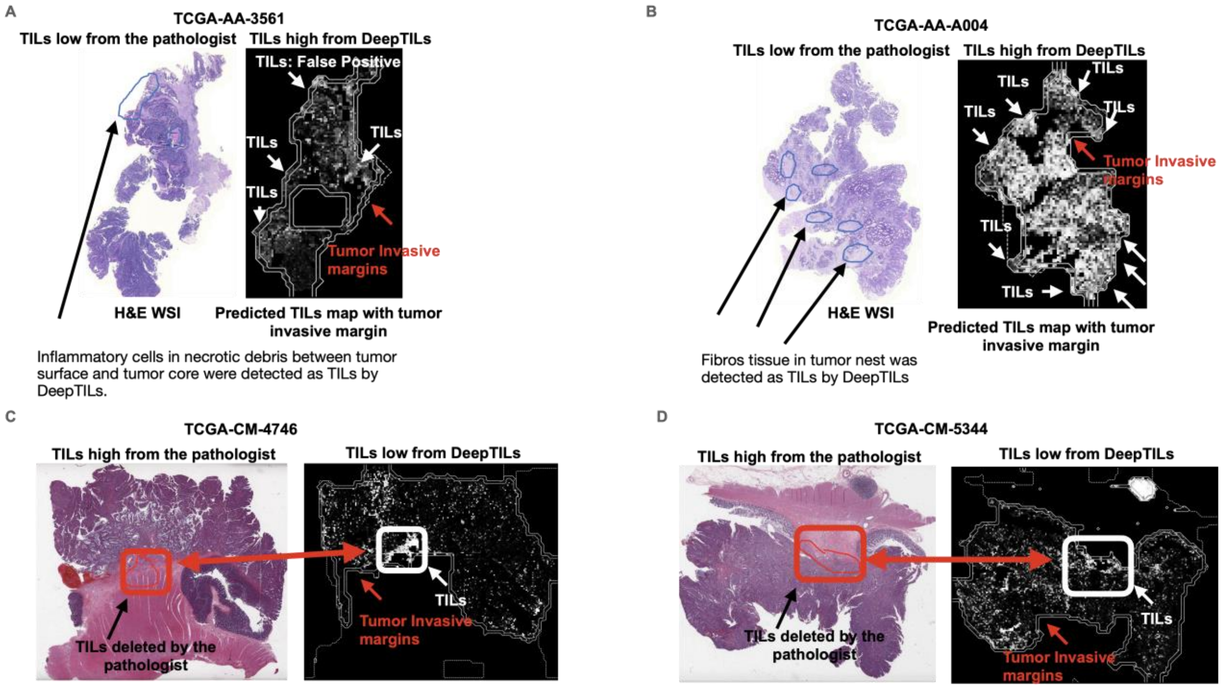


**Supplementary Figure 10. TCGA CRC patients who assigned different TILs subgroup by KM grading and DeepTILs.**

**:** (A) The pathologist graded TILs low using KM grading but DeepTILs assigned TILs high. Top left blue region indicates inflammatory cells in necrotic debris which DeepTILs identified as TILs. (B) DeepTILs identified many TILs regions but the pathologist assigned the patients as TILs low. Blue regions are fibrotic tissue regions identified by the pathologist but DeepTILs identified as TILs. Higher number of TILs detected by DeepTILs present in tumor invasive margin. (C and D) The pathologist assigned the patient as TILs high subgroup. The pathologist confirmed that most of TILs regions detected by DeepTILs are correct. Since few TILs present in tumor invasive margin detected by DeepTILs, DeepTILs assigned the patient into TILs low subgroup.
